# Inhibition of PTEN activity promotes IB4-positive sensory neuronal axon growth

**DOI:** 10.1101/2020.04.02.019042

**Authors:** Li-Yu Zhou, Feng Han, Shi-Bin Qi, Jin-Jin Ma, Yan-Xia Ma, Hong-Cheng Zhang, Ji-Le Xie, Xin-Ya Fu, Jian-Quan Chen, Bin Li, Hui-Lin Yang, Feng Zhou, Saijilafu

## Abstract

Traumatic nerve injuries have become a common clinical problem, and axon regeneration is a critical process in the successful functional recovery of the injured nervous system. In this study, we found that peripheral axotomy reduce total PTEN expression in adult sensory neurons, however, it did not alter the expression level of PTEN in IB4-positive sensory neurons. Additionally, our results indicate that the artificial inhibition of PTEN markedly promotes adult sensory axon regeneration, including IB4-positive neuronal axon growth. Thus, our results provide strong evidence that PTEN is a prominent repressor of adult sensory axon regeneration, especially in IB4-positive neurons.

## Introduction

With the rapid rise in traffic accidents, sports participation, and construction work, the incidence of nerve injury is increasing each year. However, functional recovery from nerve injury is often unsatisfactory as many patients suffer from permanent complications, such as paralysis. In the nervous system, axons are the main transmission pathway of electric impulses between two nerve cells, or between a nerve cell and a downstream effector cell. Thus, axon regeneration is a critical process for the successful functional recovery of the injured nervous system. However, there are two major factors that inhibit axonal regeneration. The first is the inhibitory external environment, including mechanical barriers like a fibrosis scar. The second is the diminished axon regeneration ability of mature neurons (Liu et al., 2010; Park et al., 2010). Previous literature has demonstrated that the manipulation of the inhibitory microenvironment can promote axonal regeneration, but the effects are limited (Lee et al., 2010). Therefore, today, enhancing the intrinsic axon regeneration ability of mature neurons has become a new strategy to promote the growth of injured axons. To date, recent studies have identified that the PTEN gene is a critical regulator of the intrinsic axon regeneration ability of mature neurons, and that PTEN knockout significantly promotes axon generation in the central nervous system (CNS) (Du et al., 2015; Kurimoto et al., 2010; Liu et al., 2010; Ning et al., 2010; Ohtake et al., 2015; Park et al., 2010).

It has been reported that mammalian dorsal root ganglia (DRG) contain several subpopulations of neurons that possess different characteristics. For example, Griffonia simplicifolia isolectin B4 (IB4) can recognize the subpopulation of neurons that expresses the receptors for glial cell line-derived neurotrophic factor (Bennett et al., 1998; Tucker et al., 2006). Additionally, IB4-labeled neurons are difficult to regenerate axon after injury, and even the forced overexpression of axon growth promoting molecules, such as α7β1-integrin and GAP43, failed to enhance the axon growth of IB4-labeled neurons (Leclere et al., 2007). These findings indicate that IB4-labeled DRG neurons lack the intrinsic ability to regenerate axons following injury. Interestingly, a recent study found that IB4-labeled neurons usually have an elevated expression of PTEN protein (Christie et al., 2010). Therefore, we hypothesized that the PTEN gene is one of the crucial inhibitors of IB4-positive (IB4^+^) neuronal axon regeneration. Thus, the suppression of PTEN activity may increase the intrinsic axon growth ability of IB4^+^ neurons, and further promote its axon regeneration after injury.

In the present study, we found that the expression of total PTEN in adult sensory neurons is down-regulated by peripheral axon injury, and inhibition of PTEN promotes its axon regeneration. However, we also found that IB4^+^ neurons possess high level of PTEN, and peripheral axotomy did not alter the expression level of PTEN in IB4^+^ neurons. Our results further indicate that artificial inhibition of PTEN markedly promotes IB4^+^ neuronal axon growth in vitro. Thus, here we provide strong evidence that PTEN is a prominent repressor of adult sensory axon regeneration, especially in IB4^+^ neurons.

## Methods and Materials

### Animals and surgical procedures

All animals (adult ICR mice, PTEN^flox/flox^ mice, and Advilin-Cre mice, from 8-10 weeks of age) were handled according to the guidelines of the Institutional Animal Care and Use Committee of Soochow University. Advilin-Cre and PTEN^flox/flox^ mice were bred together to generate DRG sensory neuronal specific PTEN knockout mice. The PTEN^flox/flox^ mice were obtained from Prof Liu Yaobo (Soochow University), Advilin-Cre mice were obtained from Zhou Fengquan (Johns Hopkins University). Surgical procedures were performed under intraperitoneal anesthesia with ketamine (100 mg/kg) and xylazine (10 mg/kg). To create the sciatic nerve injury, the skin was disinfected, and scissors were used to make a longitudinal incision of approximately 1cm. The muscles were separated, and the sciatic nerve was exposed. The sciatic nerve was then transected, and the skin incision was closed. Mice that received a sham surgery were used as controls.

### Reagents and antibodies

The neuron-specific class III β-tubulin mouse mAb (Tuj1, 1:1500) antibody was purchased from Covance. The PTEN antibody was purchased from Santa Cruz Biotechnology. Secondary antibodies conjugated with Alexa fluorophores 488 or 568 were directed against the IgGs of the primary antibody species (1:1000; Invitrogen). The Alexa Fluor 488-conjugated Isolectin IB4 was from life technology. The SF1670 and BPV were purchased from Selleck. The PTEN primer (forward: 5’-CTCCTCTACTCCATTCTTCCC-3’; reverse: 5’-ACTCCCACCAATGAACAAAC-3’) was from Gene Pharma (GenePharma Co.; Shanghai, China).

### Adult sensory neuronal culture

The Lumber 4 (L4) and L5 DRG were dissected out and treated with 2 ml 0.1% collagenase for 90 min at 37.0 °C. Then, the samples were treated again with 1 ml Tryple Express for 20 min at 37.0 °C. Tryple Express was neutralized with culture medium (the Minimum Essential Medium supplemented with 5% fetal bovine serum, 1×Penicillin/Streptomycin solution). Then, the DRGs were dissociated with a 1000 μl pipette tip using the culture medium. The dissociated cells were cultured on poly-D-lysine (100 μg/ml, Sigma) and laminin (10 μg/ml, Sigma) coated coverslips in 24-well plates. The isolated neurons were allowed to grow axons for 3 days at 37°C in a CO_2_-humidified incubator. During cell culture, we administered SF1670 (1 nM or 10 nM), or 200 nM BPV to the culture medium. As a control, the same volume of DMSO was added to the culture medium.

### Axon length analysis

All images were photographed using AxioVision 4.7 software (Carl Zeiss MicroImaging, Inc.). The longest axons of one hundred randomly selected neurons in each experimental condition were traced and measured manually using the “measure/curve” application of AxioVision 4.7 software. Average axon lengths were calculated from three separate experiments in each condition.

### Immunofluorescence staining

The samples were fixed in 4% PFA, and blocked with 2% BSA. DRG tissues were cut into slices of 14 microns and pasted on gelatin-coated glass slides. The primary antibody was applied for 24 hour at 4°C, then washed with 3% TPBS and incubated with the secondary antibodies for 1 hour at room temperature. Afterwards, tissue sections were incubated with Hoechst (1:2000) for 20 min at room temperature. For immunofluorescence staining of IB4, Alexa Fluor 488-conjugated Isolectin IB4 was applied overnight at 4°C, and then washed with 3% TPBS and mounted with a custom-made MOWIOL mounting media.

### Western blots

The proteins were extracted with RIPA buffer, and the protein concentrations were measured using Bio-Rad DC Protein Assay. The protein samples were loaded into SDS-PAGE gels, and then transferred to polyvinylidene fluoride membranes (PVDF). The PVDF membranes were incubated with the corresponding primary antibodies overnight at 4°C, and then with the secondary antibodies for 2 hours at room temperature. Protein bands were visualized using Pierce™ ECL Western Blotting Substrate, and the densities of protein bands from three independent experiments were quantified using Image J software (NIH, Bethesda, MD, USA).

### qRT-PCR

Trizol reagent was used to extract RNA from the DRG tissue. RNA was treated with reverse transcription kits and processed for cDNA synthesis. PCR primers were purchased from Sangon Biotech. The PCR products were labeled using the CFX96™ real-time PCR detection system (Bio-Rad). We used 18S as a housekeeping gene.

The following primer sequences were used:

PTEN-F: 5’-CTCCTCTACTCCATTCTTCCC-3’

PTEN-R: 5’-ACTCCCACCAATGAACAAAC-3’

18S-F: 5-TCCCTAGTGATCCCCGAGAAGT-3’

18S-R: 5-CCCTTAATGGCAGTGATAGCGA-3’

### In vivo electroporation of adult DRG neurons

Under deep anesthesia, the left L4-L5 DRGs of adult PTEN knockout mice were exposed, and 1.0μl of EGFP plasmid was micro-injected into the DRG tissue. The PTEN^flox/flox^ mice were used as the control group. After microinjection, the DRG was electroporated using a tweezer-like electrode (Ø1.0 mm) and an ECM830 Porator for five pulses (35 V, 15 ms duration, 950 ms interval). The animals were then allowed to recover. Two days later, the ipsilateral sciatic nerve was exposed, and the crush injury was made using #5 fine forceps. An 11-0 nylon suture was used to mark the crush site. After another three days, the animals were perfused transcardially with 4% PFA, and the entire sciatic nerve was dissected and removed. The lengths of all EGFP-positive axons were measured from the marking suture to the distal axon tip.

### Statistics

Data are presented as mean ± S.E.M. Two -tailed Student’s t-tests were used to compare the different experimental conditions. *P*-values of *P* < 0.05 were considered statistically significant (* *P* < 0.05; ** *P* < 0.01; *** *P* < 0.001).

## Results

### Decreased expression of PTEN in adult sensory neurons following sciatic nerve transection

Firstly, we found that, after sciatic nerve transection, the expression of total PTEN in L4/5 DRG neurons was significantly down-regulated compared to the control (Fig. 1a, b). Secondly, the PTEN mRNA level in the DRG neurons also decreased following the sciatic nerve injury (Fig. 1c), and this finding was consistent with the observed changes in protein levels. It has been reported that PTEN is an important inhibitor of the intrinsic axon growth ability of mature CNS neurons. Furthermore, the genetic knockout of the PTEN gene can significantly promote axon regeneration in the CNS (Bhagat et al., 2014; Danilov and Steward, 2015). However, the precise functional role of PTEN in the process of peripheral nervous system axon regeneration is still unknown. Using an in vitro adult peripheral sensory neuronal culture model, we found that pharmacological inhibition of PTEN activity promoted the axon growth of peripheral sensory neurons (Fig. 2a, b). Inhibition of PTEN markedly increased phospho-S6 expression in the cultured DRG neurons (Fig. 2c). Phospho-S6 is a well-known downstream target of the PTEN pathway. This result indicates that our drug concentration is enough to inhibit PTEN activity in the cultured DRG neurons. Considering the side effects of small chemical molecules, we used PTEN knockout mice to investigate their functional roles on peripheral axon regeneration. We consistently found that the axon length in the PTEN knockout mice was significantly longer than in the wild-type mice (Fig. 2d, e). Furthermore, the genetic knockout of PTEN also markedly promoted in vivo sciatic nerve axon regeneration (Fig. 2f, g). Taken together, our results indicate that the inhibition of PTEN activity promotes axon growth in adult sensory neurons in vitro and in vivo.

**Figure 1.**
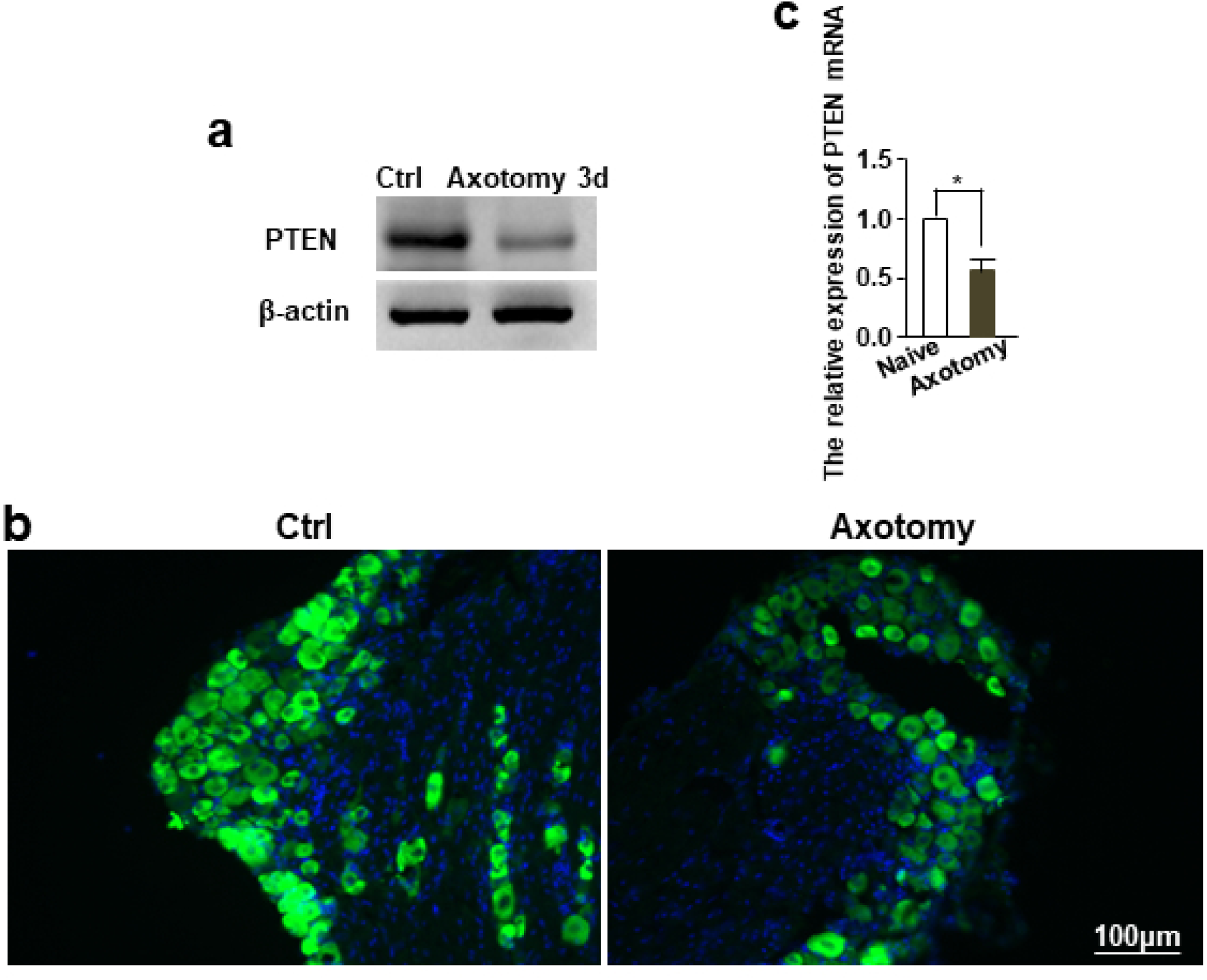
Injury induces down regulation of PTEN expression in adult sensory neurons. (a) Representative Western Blot images of PTEN expression in L4-L5 DRGs of adult mice 3 days after sciatic nerve axotomy. Compared to uninjured control, PTEN protein level is markedly down-regulated by peripheral axotomy (n=3). (b) Immunohistological staining showed PTEN expression was decreased in the L4 DRG tissues after sciatic nerve axotomy. Green: PTEN, Scale bar: 100 μm. (c) Quantification of PTEN mRNA levels by qPCR. The relative level of PTEN mRNA expression in the L4 DRGs was reduced 3 days after sciatic nerve axotomy (n=3). * *P* <0.05.

**Figure 2.**
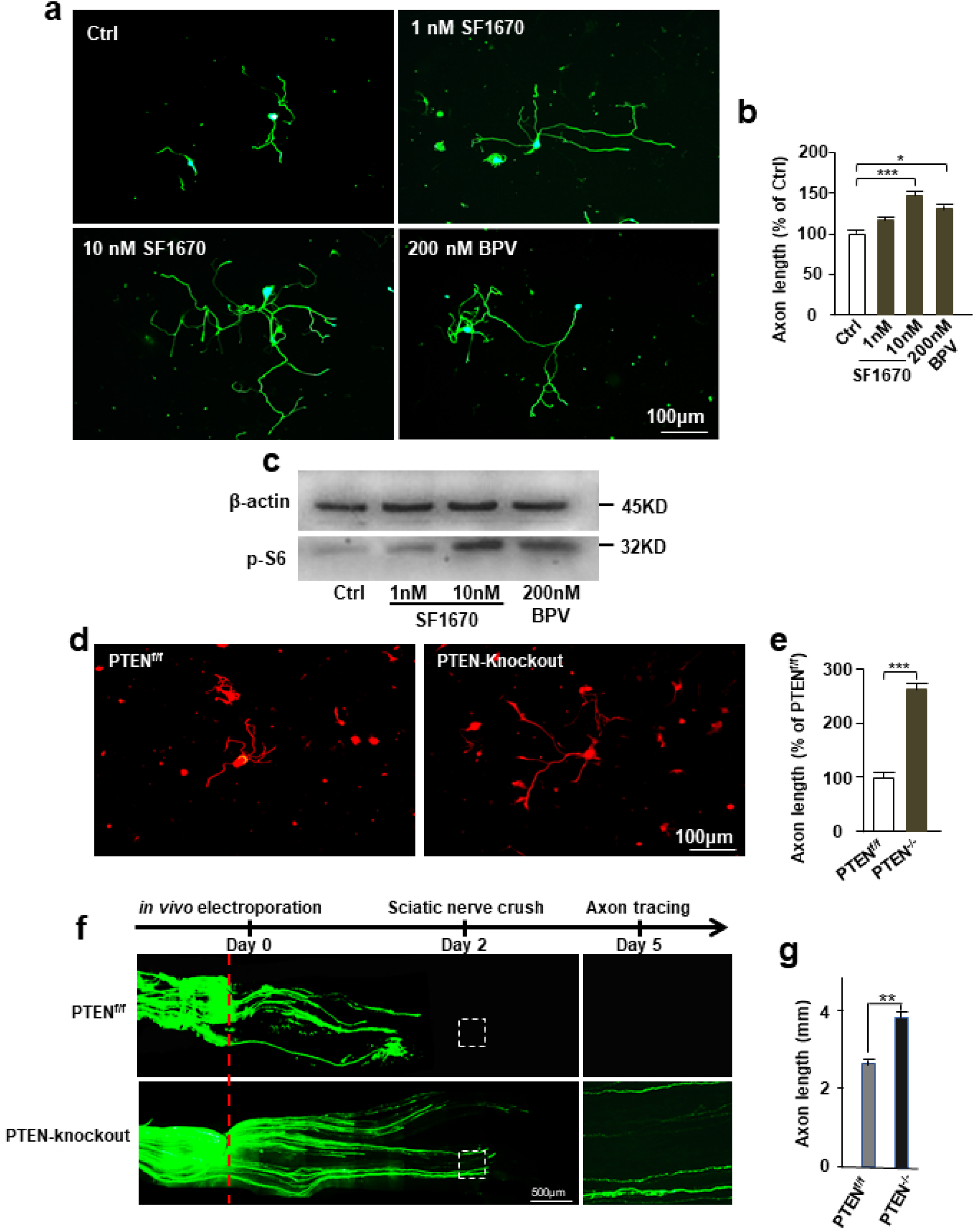
PTEN inhibition promotes sensory axon regeneration in vitro and in vivo. (a) Adult DRG neurons were cultured in vitro for 3 days, and treated with PTEN inhibitor SF1670 (1nM or 10 nM) or BPV (200 nM). The vehicle DMSO was as control. All neurons were stained with anti-βIII tubulin (green). Scale bar: 100μm. Quantification of the average length of the longest axons (n = 3). (b) The pharmacological inhibition of PTEN activity promoted the axon growth of peripheral sensory neurons (n=3). * *P* <0.05,*** *P* <0.001. (c) The pharmacological inhibition of PTEN activity markedly increases phospho-S6 expression, which is a well-known downstream target of the PTEN, in the cultured DRG neurons. (d) The adult DRG neurons were cultured from the Advilin-Cre induced PTEN knockout mice. Scale bar: 100μm. (e) Axon length of adult DRG neuron in the PTEN knockout mice was significantly longer than the PTEN^flox/flox^ mice (n=3). *** *P* <0.001. (f) The left L3-L4 DRGs were electroporated with EGFP plasmid and sciatic nerve was crushed two days after electroporation. Another three days later, whole sciatic nerve segment was harvested. Red dot line is crush site. Arrow head is regenerating axons. Scale bar: 500μm. (g) PTEN knockout significantly promotes sciatic nerve axon regeneration in vivo (n=5). *** *P* <0.001.

### IB4^+^ neurons have a higher PTEN expression level than IB4^-^ neurons,and PTEN expression was not significantly decreased by sciatic nerve transection

Adult sensory neurons can be divided into two subpopulations according to their IB4 binding ability, referred to as IB4^+^ and IB4-negative (IB4^-^) neurons. IB4^+^ neurons have very limited axon regeneration ability, and high levels of expression of PTEN protein. In agreement with previously published literature (Christie et al., 2010), our immunofluorescence staining results revealed that PTEN expression in the IB4^+^ neurons was significantly higher than in the IB4^-^ neurons (Fig. 3a). Additionally, we found that the average axon length of the IB4^+^ neurons was significantly shorter than the IB4^-^ neurons (Fig. 3b, c). Previous findings have demonstrated that peripheral axotomy can activate the intrinsic axon regeneration ability of adult sensory neurons (Smith and Skene, 1997). For this reason, we investigated the effects of peripheral axotomy on PTEN expression in IB4^+^ neurons. We found that peripheral axotomy did not affect the expression level of PTEN proteins in the IB4^+^ neurons (Fig. 4a). Furthermore, axon growth from the IB4^+^ neurons was not promoted by peripheral axotomy (Fig. 4b, c). However, we observed that peripheral axotomy significantly promote axon regeneration in IB4^-^ neurons (Fig. 4b, c). These results suggested that PTEN protein is a potential inhibitor of the axon growth ability of IB4^+^ neurons.

**Figure 3.**
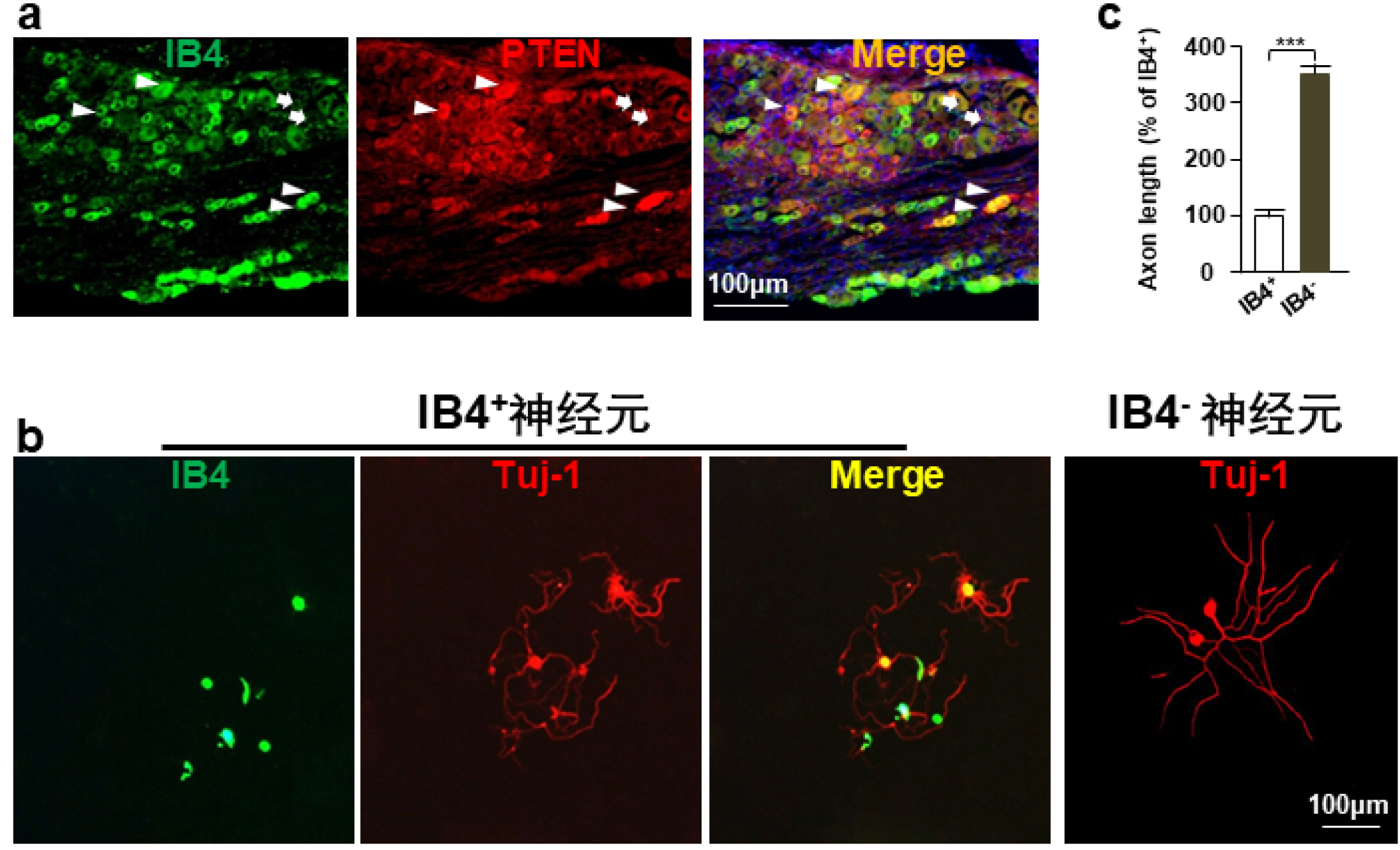
IB4^+^ neurons possess higher PTEN expression, and lower intrinsic axon growth ability. (a) Immunohistological staining of L4 DRG section showed that IB4+ neurons possess higher PTEN expression. Scale bar, 100μm. (b) Adult DRG neurons were cultured for three days, and stained with anti-βIII tubulin (red) and Alexa Fluor 488-conjugated Isolectin IB4 (green). Scale bar, 100μm. (c) Axon length from IB4^+^ neurons was significantly shorter than IB4^-^ neuron (n = 3). *** *P* <0.001.

**Figure 4.**
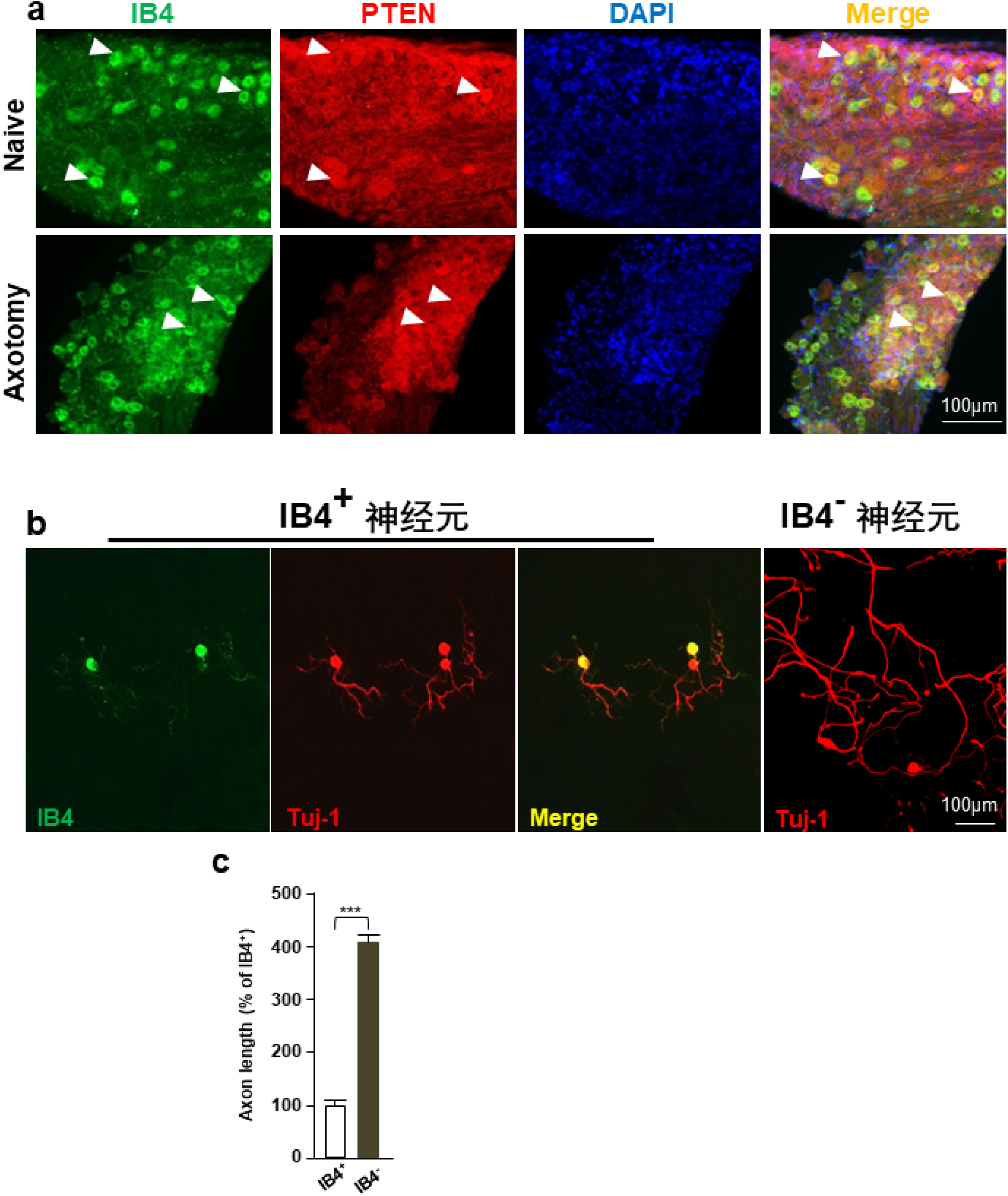
PTEN expression in IB4^+^ neurons was not affected by sciatic nerve axotomy. (a) Immunohistostaining of L4 DRG section showed that PTEN expression in IB4^+^ neurons was not affected by sciatic nerve axotomy. Arrowhead : IB4^**+**^ neurons. Scale bar, 100μm. (b) Adult DRG neurons were culture one day after peripheral nerve axotomy, and stained for anti-βIII tubulin (red) and Alexa Fluor 488-conjugated Isolectin IB4 (green). Scale bar, 100μm. (c) IB4^-^ neuronal axon growth is significantly enhanced by peripheral axotomy, however, intrinsic axon growth ability of IB4^+^ neuron is not affected (n=3). *** *P* <0.001.

### Inhibition of PTEN promotes axon regeneration in IB4^+^ sensory neurons

Next, to investigate the effect of PTEN on IB4^+^ neuronal axon regeneration, we administrated a specific PTEN inhibitor, either 10 nM SF1670 or 200 nM BPV in cultured DRG neurons (Christie et al., 2010). The pS6 is a well-known downstream substrate of PTEN, and previous findings have shown that inhibition of PTEN can promote phosphorylation of pS6. In accordance with these previous findings, we found that the level of phospho-pS6 protein significantly increased in the drug treatment group (Fig. 5a). Additionally, we also found that the axon length of IB4^+^ neurons was much longer in the SF1670 and BPV treated groups than in the control group (Fig. 5b, c). In accordance with the pharmacological results, the axon length of the IB4^+^ neurons from the PTEN knockout mice was also significant increased compared to the control mice (Fig 6a, b).

**Figure 5.**
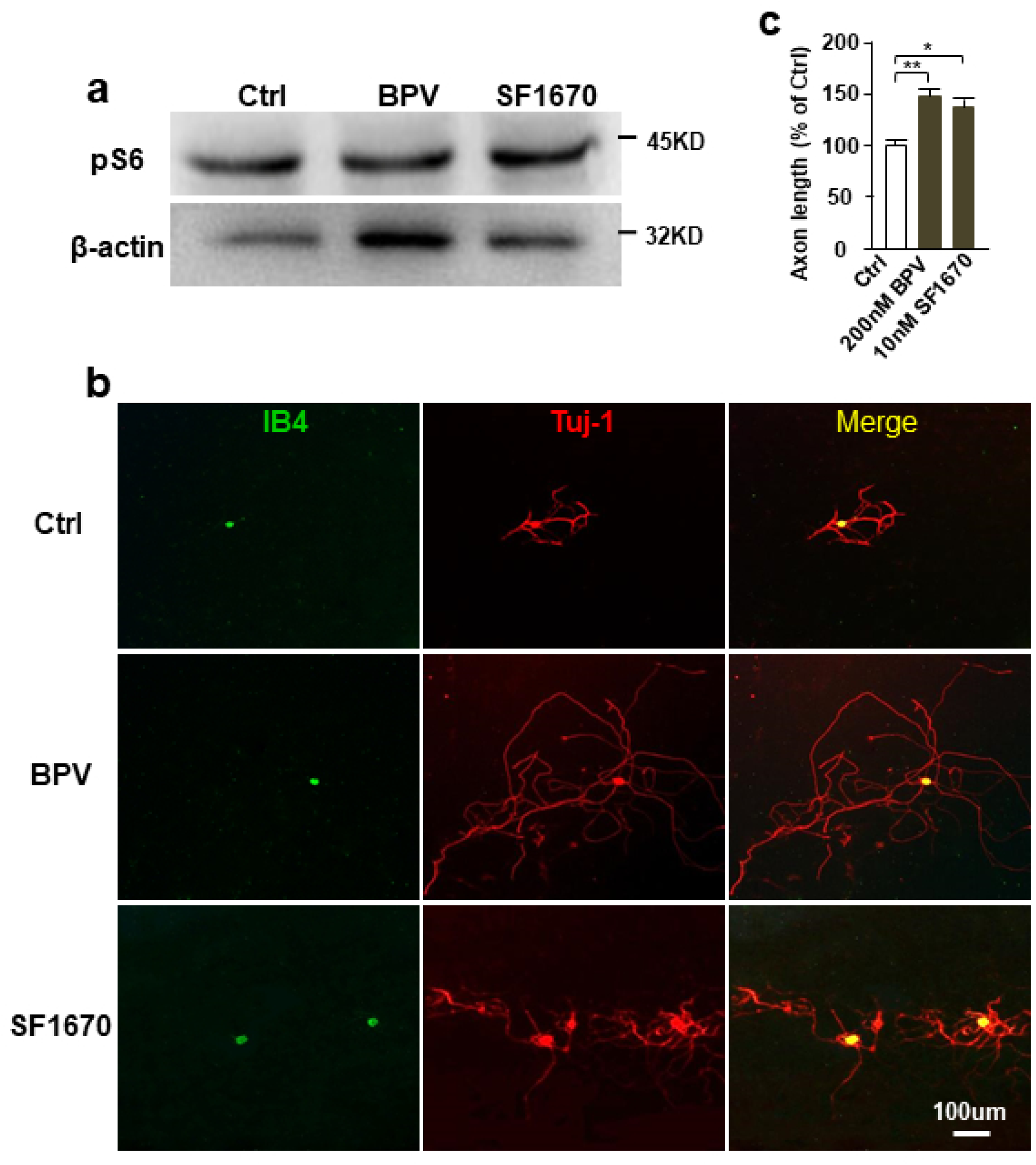
PTEN inhibition dramatically promotes IB4^+^ neuronal axon growth. (a) Representative western blot images of phosphor-S6 in adult DRGs neuronal culture with PTEN inhibitor SF1670 or BPV, and vehicle DMSO were as control group. (b) Adult DRG neurons were cultured for three days with PTEN inhibitor SF1670 (10 nM) or BPV (200 nM) treatment. Vehicle (DMSO) was as control. All neurons were stained with anti-βIII tubulin (red) and Alexa Fluor 488-conjugated Isolectin IB4 (green). Scale bar, 100μm. (c) Administration of PTEN inhibitor significantly increases axon length of IB4+ neuron (n=3). * *P* <0.05,** *P* <0.01.

**Figure 6.**
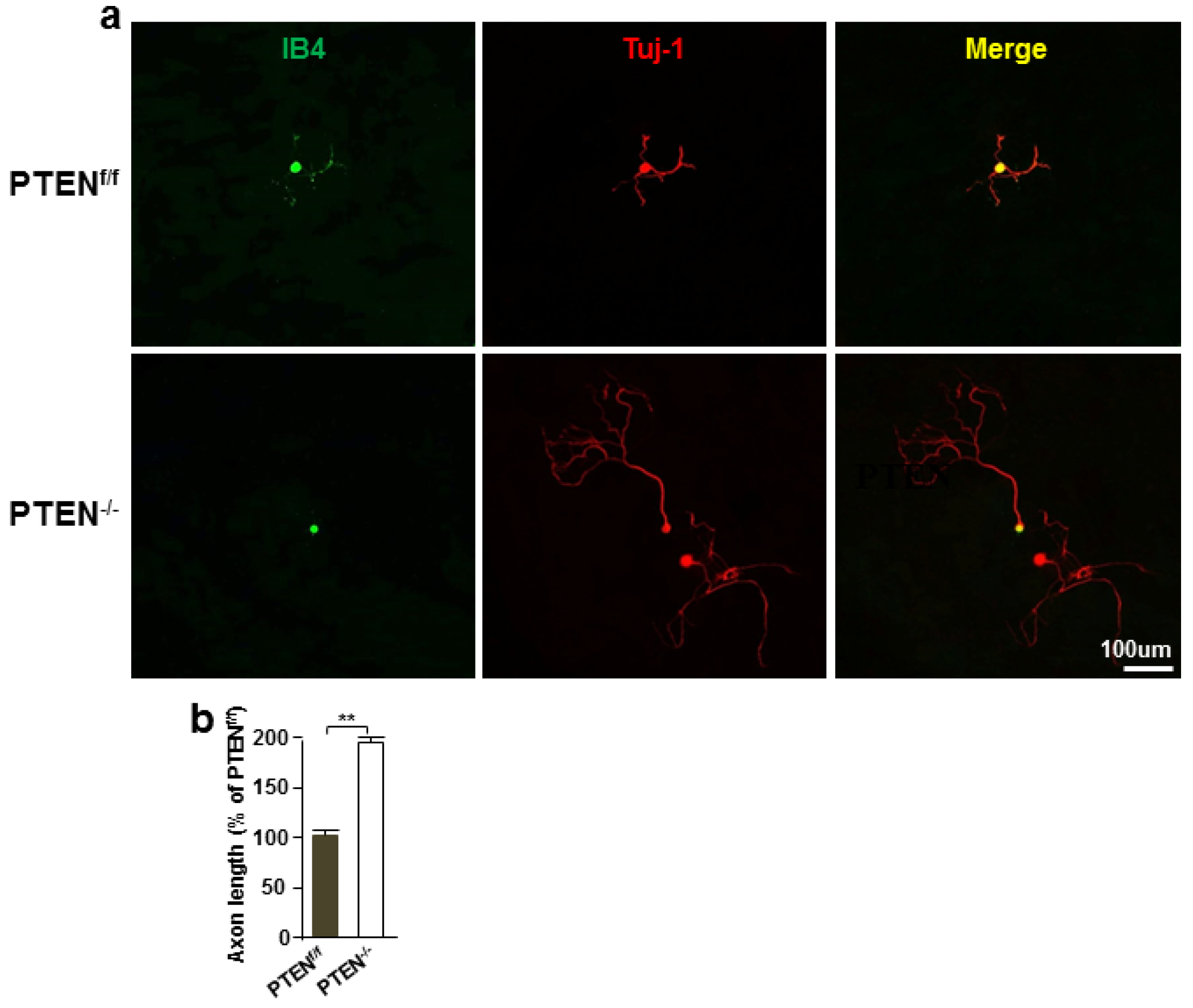
Knockout of PTEN gene promotes IB4^+^ neuronal axon growth. (a) Adult DRG sensory neurons were culture for three days from advilin-cre induced PTEN knockout mice. All neurons were stained with anti-βIII tubulin (red) and Alexa Fluor 488-conjugated Isolectin IB4 (green). Scale bar, 100μm. (b) Quantification of the average axon length showed that PTEN knockout markedly promotes IB4^+^ neuronal axon growth (n=3). ** *P* <0.01.

### PTEN inhibition of peripheral axon growth is independent of the mTOR pathway

It has been reported that the activation of the mTOR pathway can enhance axon outgrowth of adult CNS axon regeneration (Lim et al., 2016; Park et al., 2008), and that mTOR activity is regulated by PTEN. Therefore, to determine whether PTEN regulates sensory axon growth via mTOR activity, we simultaneously blocked both PTEN and mTOR activity using a specific pharmacological inhibitor. The adult DRG neurons were cultured for 3 days with both PTEN and mTOR inhibitors, and axon length was measured. Our data indicated that mTOR inhibition did not affect the axon growth promoting effect of the PTEN inhibitor (Fig 7a, b). There was no obvious difference in axon length between the group treated with PTEN inhibitor only and the group treated with PTEN inhibitor combined with rapamycin. This means that the regulatory effect of PTEN on adult peripheral sensory neurons is not dependent on the mTOR pathway.

**Figure 7.**
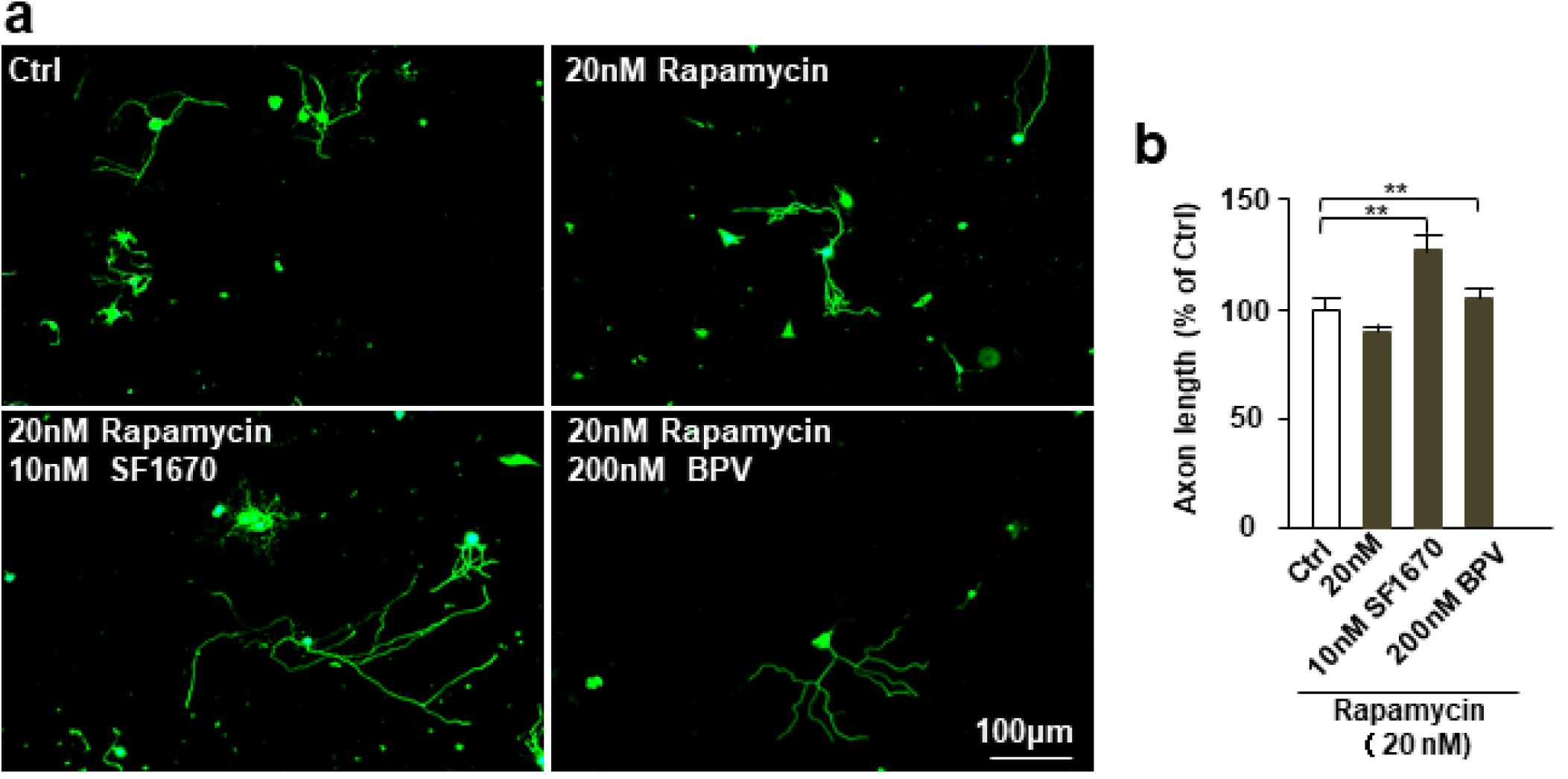
PTEN inhibition induced peripheral axon growth is independent of the mTOR pathway. (a) Adult DRG neurons were cultured with rapamycin (20nM) only, or cultured with both rapamycin 20nM and PTEN inhibitor (10nM SF1670 or 200nM BPV). All neurons were stained with anti-βIII tubulin (green). Scale bar, 100μm. (b) Quantification of the average axon length showed that mTOR inhibition did not affect the axon growth promoting effect of the PTEN inhibitor (n=3). ** *P* <0.01.

## Discussion

Peripheral nerve injuries often result in the loss of related sensory and motor functions, and it has been hypothesized that damaged peripheral sensory neurons can spontaneously regenerate axons. However, recent accumulating evidence has shown that only some parts of adult sensory neurons can regenerate axons after injury. Researchers have characterized multiple subgroups of adult DRG neurons according to their morphological and phenotypic differences. For example, based on their lectin Griffonia simplicifolia IB4 binding ability, adult DRG neurons can be classified as either IB4^+^ or IB4^-^. Out of these two types, IB4^+^ neurons have neither p75 neurotrophin receptors, nor Tyrosine kinases (Trk) (Christie et al., 2010). Therefore, IB4^+^ neurons possess very limited axon regeneration abilities. It has been well established that a primary injury of adult sensory neurons, known as a ‘conditioning lesion’, can prime sensory neurons into an active growth state, and dramatically increase their intrinsic axon regenerative ability. However, even a conditioning lesion cannot promote axon regeneration of IB4^+^ neurons.

The specific phosphatase activity of tumor suppressor gene PTEN (Guo et al., 2002; Li et al., 1997) was first discovered in 1997. PTEN can dephosphorylate phosphatidylinositol-3,4,5-diphosphate(PIP3)togenerate phosphatidylinostitol-4,5-diphosphate (PIP2). By doing so, PTEN regulates the intrinsic axon growth ability of adult retina ganglion cells (RGCs) and cortical neurons, and therefore plays a predominant role in mammalian nerve regeneration (Liu et al., 2010; Park et al., 2008). Previous findings have shown that knocking out the PTEN gene significantly promotes CNS axon regeneration in the optic nerve and in the cortical spinal tract. Additionally, PTEN gene expression increases during embryonic development, and, at the same time, the axon regeneration ability of mammalian neurons decreases with embryonic maturation. These studies clearly indicate that PTEN is a critical inhibitor of axon regeneration. Interestingly, it has been reported that IB4 sensory neurons possess high levels of PTEN proteins (Christie et al., 2010). In agreement with these previous findings, our immunohistological staining also revealed that the expression of PTEN in IB4^+^ neurons was significantly higher than in IB4^-^ neurons. Most adult animal DRG neurons express laminin receptor alpha7beta1 (Ekstrom et al., 2003), and its expression increases after peripheral nerve injury (Wallquist et al., 2004; Werner et al., 2000), thereby promoting axon growth on laminin (Lankford et al., 1998; Smith and Skene, 1997). However, Gardiner, et al (Gardiner et al., 2005) found that IB4^+^ neurons lack integrin alpha 7 and GAP43, which is a marker of axon regeneration (Benowitz and Routtenberg, 1997). Enhanced overexpression of both integrin alpha 7 and GAP43 cannot promote IB4^+^ neuronal axon regeneration. However, our results showed that conditioning lesion does not alter PTEN expression. Our data also showed that the inhibition of PTEN activity with its specific pharmacological inhibitor markedly promoted IB4 neuronal axon regeneration. Furthermore, knocking out PTEN in adult sensory neurons also significantly enhances axon regeneration in vitro and in vivo, and these findings are in accordance with a previous study on CNS axon regeneration. Thus, our data clearly indicate that PTEN is a critical repressor of IB4^+^ neuronal axon regeneration.

It has been reported that PTEN regulates mature CNS axon regeneration through the activation of the mammalian target of rapamycin (mTOR) pathway (Liu et al., 2010; Park et al., 2008). Our data confirmed that the deletion of PTEN can also elevate the axon regenerative ability of adult peripheral sensory neurons. Additionally, the specific pharmacological inhibition of PTEN significantly promoted axon growth in the cultured DRG neurons. However, the mTOR inhibitor did not affect the axon growth promoting effect of PTEN deletion. Additionally, the mTOR inhibitor alone did not influence axon growth of the adult sensory neurons. Therefore, the results suggest that PTEN regulates sensory neuronal axon growth independently of the mTOR pathway. PTEN also regulates Glycogen synthase kinase-3 beta (GSK3β) activity independently of the mTOR pathway. Phosphorylation of GSK3β by PTEN-Akt is known to be the major mechanism by which GSK3β is inactivated. An earlier study conducted by our team showed that the inactivation of GSK3β is necessary for sensory axon regeneration (Saijilafu et al., 2013). Thus, it is possible that deletion of PTEN promotes sensory axon regeneration through GSK3β activity.

In conclusion, our results demonstrated that IB4^+^ sensory neurons express a high level of PTEN, and that even peripheral axotomy does not modify PTEN expression in IB4^+^ sensory neurons. Furthermore, we found that PTEN inhibition dramatically promotes axon growth of both IB4^+^ sensory neurons and IB4^-^ neurons. Taken together, these data indicate that PTEN is a key intrinsic regulator of axon growth ability of adult sensory neurons.

## Author Contributions

LY Z, F H, SB Q, JJ M, YX M, HC Z, JL X, XY F, JQ C, B L, HL Y, F Z, and Saijilafu designed the experiment. LY Z, F H, and SB Q performed the experiments. LY Z, F H, and Saijilafu co-wrote the manuscript.

## Conflict of Interest Statement

The authors declare that there is no competing interest.

## Data Accessibility Statement

The data that support the findings of this study are available from the corresponding author upon reasonable request.

## Notes

**Funding Statement**: This work was supported by a grant from the National Natural Science Foundation of China (No. 81571189 and No. 81772353 to Saijilafu), National Key Research and Development Program (No. 2016YFC1100203), An Innovation and Entrepreneurship Program of Jiangsu Province, and A Priority Academic Program Development of Jiangsu Higher Education Institutions.

